# MinION Sequencing of colorectal cancer tumour microbiomes – a comparison with amplicon-based and RNA-Sequencing

**DOI:** 10.1101/662270

**Authors:** William S Taylor, John Pearson, Allison Miller, Sebastian Schmeier, Frank A Frizelle, Rachel V Purcell

**Affiliations:** Department of Surgery, University of Otago, Christchurch, New Zealand; Biostatistics and Computational Biology Unit, University of Otago, Christchurch, New Zealand; Gene Structure and Function Laboratory, University of Otago, Christchurch, New Zealand; Institute of Natural and Mathematical Sciences, Massey University, Auckland, New Zealand

## Abstract

Recent evidence suggests a role for the gut microbiome in the development and progression of colorectal cancer. In this study we compare MinION sequencing with meta-transcriptomics and amplicon-based sequencing for microbiome analysis of colorectal tumour tissue samples. DNA and RNA were extracted from 11 colorectal tumour samples. RNA-Sequencing, 16S rRNA amplicon sequencing and MinION genomic sequencing was carried out and resulting data used as input for taxonomic classification using Kraken2. Taxonomic concordance between the three platforms at different taxonomic levels was tested on a per sample basis. The average number of reads per sample using RNA-Sequencing was more than 129 times that generated using MinION sequencing. However, the average read length of MinION sequences was more than 13 times that of RNA or 16S rRNA amplicon sequencing. 16S sequencing was less informative beyond the genus level, and both RNA-Sequencing and MinION sequencing could detect more phyla and genera in the same samples, compared to 16S sequencing. Long-read sequences generated using MinION sequencing can compensate for low numbers of reads for bacterial classification. MinION sequencing can discriminate between bacterial strains and plasmids and shows potential as a tool for microbiome sequencing of colorectal cancers in a clinical setting.

## Introduction

Colorectal cancer is associated with high mortality and is becoming increasingly common [1, 2]. The best method of prevention is early detection through mass screening programs, such as faecal testing and sigmoidoscopy of at-risk individuals [3]. However, these are not adapted for early-stage tumours. Lifestyle and diet are the strongest risk factors for the disease, and changes in the gut microbiome have been linked to colorectal cancer development and pathogenesis in an increasing number of studies [4]. The assessment of microbiomes in a clinical setting is not widely practiced other than screening faecal material for individual pathogens such as *Helicobacter pylori* [5]. Reasons for this include the lack of access to sequencing technology and clinical training for the interpretation of microbiome sequencing data [6]. Sampling, library preparation, and sequencing are expensive and time consuming, which makes it less feasible for many applications [7, 8].

Metagenomic samples are typically assessed by amplicon sequencing using marker genes; the bacterial 16S ribosomal RNA (rRNA) gene is the most commonly used. This gene is highly conserved in bacteria and contains variable regions that can be used for taxonomic differentiation [9]. 16S rRNA sequencing is the current “gold standard” for bacterial classification, and protocols and downstream analysis tools are widely available [10]. Amplicon-based sequencing can be reliably used to detect genus- and phylum-level differences, but has less power for detecting species- and strain-level differences [11]. Additionally, the technique lacks the ability to analyse plasmids or any genomic region outside of the marker gene, e.g. virulence or antibiotic resistance genes, or variations unique to a species/strain [12, 13].

Meta-transcriptomics using RNA-Sequencing is a powerful tool for interrogating transcribed genes in a sample [14] and can be utilised for microbial classification based on specific RNA transcripts [15].

However, RNA-sequencing is a labour-intensive task requiring costly reagents and specialised protocols. Currently, bioinformatics of meta-transcriptomics data requires specifically designed software, which is more computationally intensive, in comparison to tools for human genomic data analysis [16].

Oxford Nanopore Technologies have developed a small, inexpensive and portable sequencing platform, the MinION, which addresses many of the shortcomings of other available typical next-generation sequencing platforms, such as cost, reagent usage, and analysis bottlenecks, by utilising a non-synthesizing sequencing method [8, 17]. This platform can produce sequencing reads from all genomic material in an environmental or clinical sample and allows taxonomic classification of elements that may not be detected using marker genes or actively transcribed gene sequencing alone.

To investigate the utility of MinION sequencing for complex microbiome analysis, this study has compared the microbiome component of 11 colorectal cancer tissue samples using three different sequencing methods: RNA-Sequencing, 16S rRNA amplicon sequencing, and MinION sequencing. The study used Kraken2 for rapid k-mer based assignment of taxa. The aim was to determine the applicability of using MinION sequencing and a single flow-cell as a method for rapid, cost-effective bacterial taxonomic classification of clinical tissue samples.

## Methods

### Patient cohort

Colorectal cancer samples were obtained from treatment-naïve tumours during surgical resection. Study participants gave informed written consent, and all experimental protocols were approved and carried out in compliance with the University of Otago Human Ethics Committee regulations (ethics approval number: H16/037).

### DNA and RNA extraction

Tissue samples were frozen in liquid nitrogen and stored at −80°C post resection, and subsequently transferred to RNAlater ICE^TM^ (Qiagen) and stored at −20°C. As described previously [18], nucleic acid extraction was performed on < 20mg of tissue by a single operator in one batch to avoid variation in protocol. Tissue disruption was carried out using a Retch Mixer mill. DNeasy Blood and Tissue Mini Kit (Qiagen) and RNEasy Plus Mini Kit (Qiagen) were used for DNA and RNA extraction, respectively. Quantification of the extracted nucleic acids was carried out using a NanoDrop 2000c spectrophotometer (Thermo Scientific, Asheville, NC, USA) and samples were stored at −80°C.

### 16S rRNA sequencing and RNA sequencing

16S rRNA amplicon sequencing (16S-Seq) and RNA-Sequencing (RNA-Seq) data was accessed from publicly available sequence data stored in Sequence Read Archive, as study SRP117763 [19]. The data files corresponding to the samples accessed in the current study are given in Supplementary Table S1.

### MinION library preparation

Eleven DNA samples from colorectal cancer tissue, corresponding to those used for 16S rRNA sequencing were used for MinION genomic DNA sequencing (MinION-Seq) (see Supplementary Table S1). DNA concentration was recorded with a Qubit**®** 2.0 fluorometer prior to library preparation. A reagent blank (nuclease-free water) was included as a technical control. Size selection was performed on each of the samples to remove fragments less than ~400bp, using 0.45x the volume MagBio High Prep beads. For each sample, 400ng genomic DNA was used, the volume adjusted to 7.5µl with nuclease-free water, and 2.5µl of barcode fragmentation mix was added, as per MinION protocol RBK_9054_v2_revA. The samples were incubated in a thermal cycler at 30°C for 1 minute and 80°C for 1 minute. The 12 barcoded samples were then pooled, and DNA was purified and resuspended in 10µl of 10mM Tris-Hcl pH 7.5 with 50mM NaCl. 1µl of RAP (Rapid sequencing AdaPtor) were added to the barcoded DNA. The resulting library was loaded into a MinION R9.4.1 (106) flow cell and sequenced for 48 hours.

### Sequence processing

#### 16S rRNA Amplicon Sequencing

Data was accessed from the publicly available sequence data stored in the Sequence Read Archive as study SRP117763 [19]. In brief, the short overlapping forward and reverse reads from the same fragments were joined using FLASh v1.2.11 [20], joined overlapped sequences were trimmed to contain only those reads with a 99.99% accuracy. Minimal length of fragments was kept at 50bp using SolexaQA++ v3.1.15 [21] and chimeric sequences were removed using scripts in QIIME v1.9 [22].

#### RNA-Sequencing

A GRCh38p12 human genome index with RefSeq annotation was generated and RNA sequences mapped using STAR v2.5.3a [23]. Unmapped reads were converted to FASTQ, sorted, and separated from mapped reads using samtools [24]. Bedtools was used to extract unmapped reads as FASTQ files for use in subsequent taxonomic classification analysis [25]. Singletons and paired datasets were combined post taxonomic classification.

#### MinION genomic DNA sequencing

Base calling and first-pass demultiplexing was performed using Albacore v2.3.3. Sequence quality analysis was performed using NanoPlot 0.16.4 [26]. Porechop v0.2.3 was used to remove barcodes and adaptors, and to verify Albacore demultiplexing. Reads were filtered for quality and length, >Q7 and >120bp, respectively, using FiltLong v0.2.0. The GRCh38p12 human genome index was created using Minimap2 [27] v2.14-r883, to which all MinION reads were mapped. Unmapped reads were extracted and converted first to BAM and then to FASTQ format using samtools v1.9 [24].

### Classification

#### Taxonomic assignment

Kraken2 v2.0.6-beta was used for taxonomic classification of all datasets [28]. The genetic data for constructing the databases were retrieved from the NCBI RefSeq library. The database (ba_CRC) contained archaea and bacterial taxa complete genomes from the RefSeq NCBI database (Supplementary Table S2) and included partial assemblies of selected taxa from a group of species known to be associated with CRC (Supplementary Table S3). Results were tabulated and analysed using Pavian [29]. Integrated Genomics Viewer v2.4.18 [30] was used to analyse and visualise alignment of sequencing reads with selected bacterial genomes.

#### Statistics

Spearman correlation analysis was carried out using RStudio (R version 3.4.4).

## Results

### Read number and length of sequences for each platform

The average number of raw reads per sample was 13,951,214, 233,193 and 68,534 for RNA-Seq, 16S-Seq and MinION-Seq, respectively. After processing the raw reads (quality control and removing reads mapping to the human reference), the per sample average number of reads was 1,941,172, 141,743 and 15,020 for RNA-Seq, 16S-Seq and MinION-Seq, respectively.

RNA-Seq and 16S-Seq paired-end reads were 250bp and 125bp in length, respectively, giving a 250 and 500bp query maximum for these platforms, while the average MinION-Seq read length was 1631bp, giving a genomic query sequence more than three times longer on average; the longest MinION read, at 46,392bp, was 92 times longer than the other platforms.

### Mapping to the human genome

RNA-Seq reads were mapped to the human genome GRCh38p12 using the STAR aligner. An average of 1.9 million unmapped reads remained per sample after mapping, 13.9% of the total. MinION-seq reads were mapped to a GRCh38p12 index using MiniMap2, an average of 762 non-human reads remained per sample after mapping, 16.9% of the total. All unmapped reads were putatively bacterial and were used in subsequent classification analysis. Taxonomic profiles to the species level were generated using Kraken2 for all three sequencing platform data (Supplementary Table S4).

### Comparison of bacterial taxonomic profiles derived using different sequencing platforms

The per sample concordance between the sequencing platforms was found to be highest at the phylum level between RNA-Seq and MinION-Seq. However, at the genus and species levels, the correlation between 16S-Seq and RNA-seq was stronger (Figure 1).

**Figure 1.**
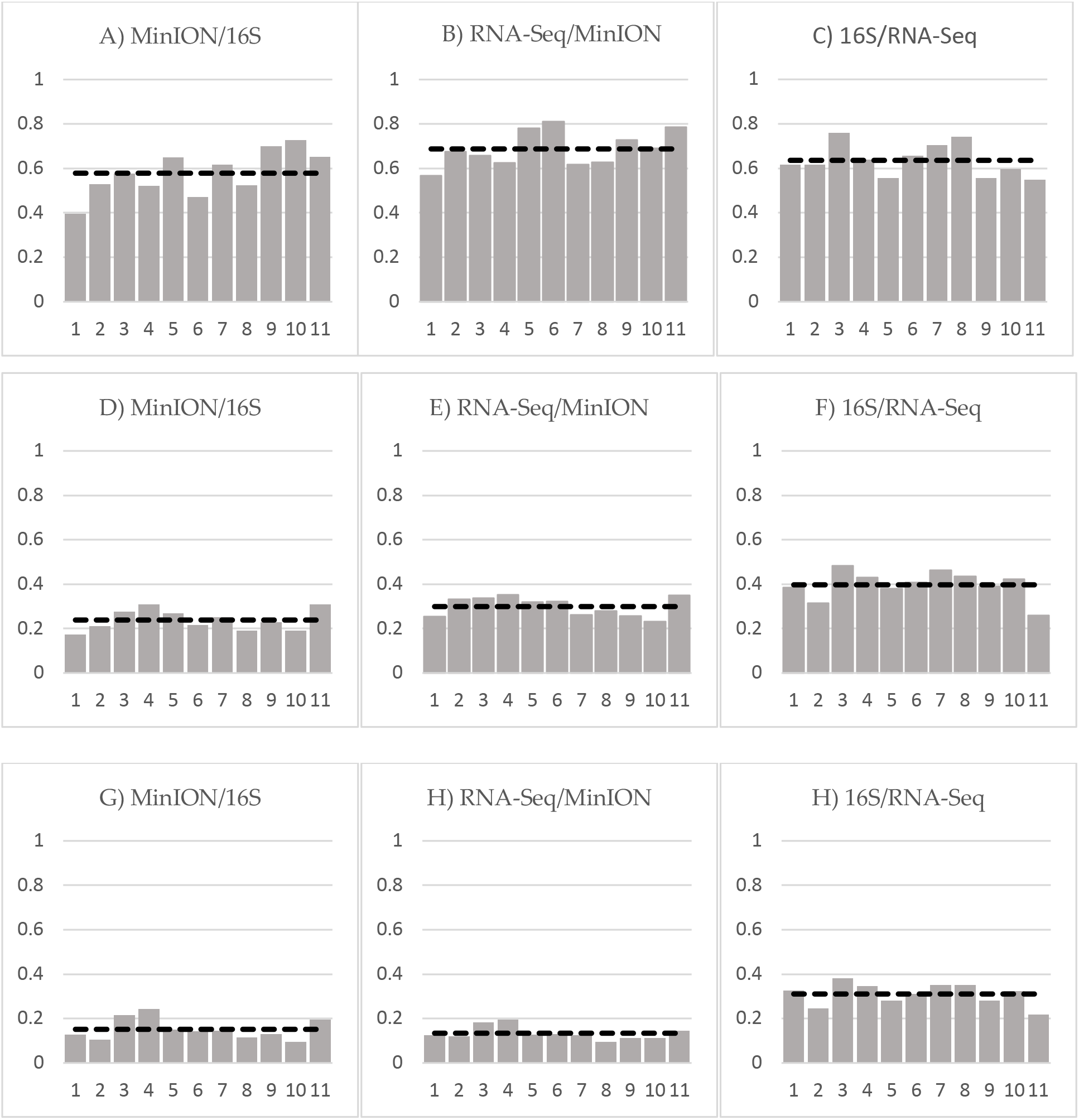
Level of concordance per sample between platforms using Spearman’s rank correlation, using all detected and absent taxa data at A–C) the phylum level, D–F) the genus level, and G–I) the species level. The dashed line indicates the average concordance across all samples.

Using Spearman’s rank correlation (Figure 1), we found an average 63.5%, 39% and 31% concordance between 16S-Seq and RNA-Seq at the phylum, genus and species levels respectively. Concordance between RNA-Seq and MinION-Seq was 68.7%, 29.9% and 13.3% at the phylum, genus and species levels, respectively, and concordance between 16S-Seq and MinION-Seq was 57.8%, 23.8% and 19.6% at the phylum, genus and species levels respectively, per sample.

The taxa identified at each taxonomic level from all 11 samples combined, was substantially different between sequencing platforms, ranging from 80.5% similarity in identified phyla to only 18.9% similarity at the species level between 16S-Seq and RNA-Seq (Table 1).

**Table 1.**
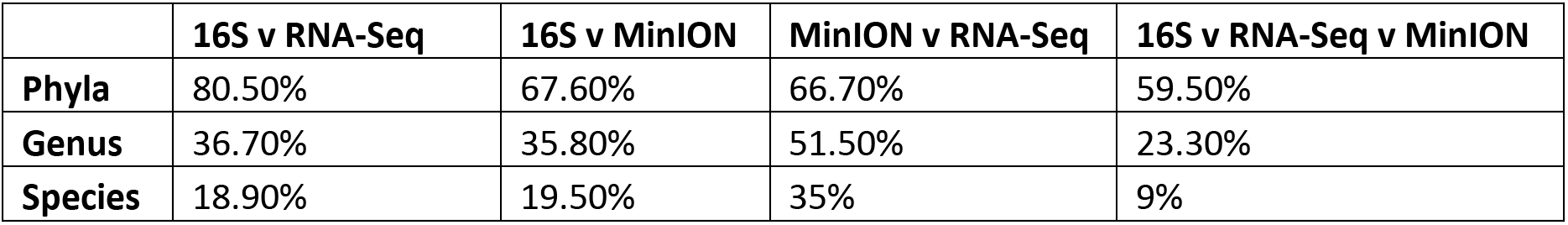
Similarity in taxa identified between platforms

### Microbiome sample composition

To estimate the relative abundance of taxa within samples using Kraken2, we used the number of assigned reads as an indicator of an individual bacteria. Figure 2A shows the proportional composition detected using each of the platforms, as estimates of abundance of different taxa. The relative abundance of phyla detected using each platform varied, with 16S-Seq detecting higher levels of Firmicutes than Bacteroidetes, compared to the other platforms, while a larger proportion of MinION sequencing reads were assigned to Proteobacteria. As expected, Firmicutes, Bacteroidetes, Fusobacterium and Proteobacteria were the most abundant phyla detected, although their relative abundance differed depending on the sequencing platform used.

**Figure 2.**
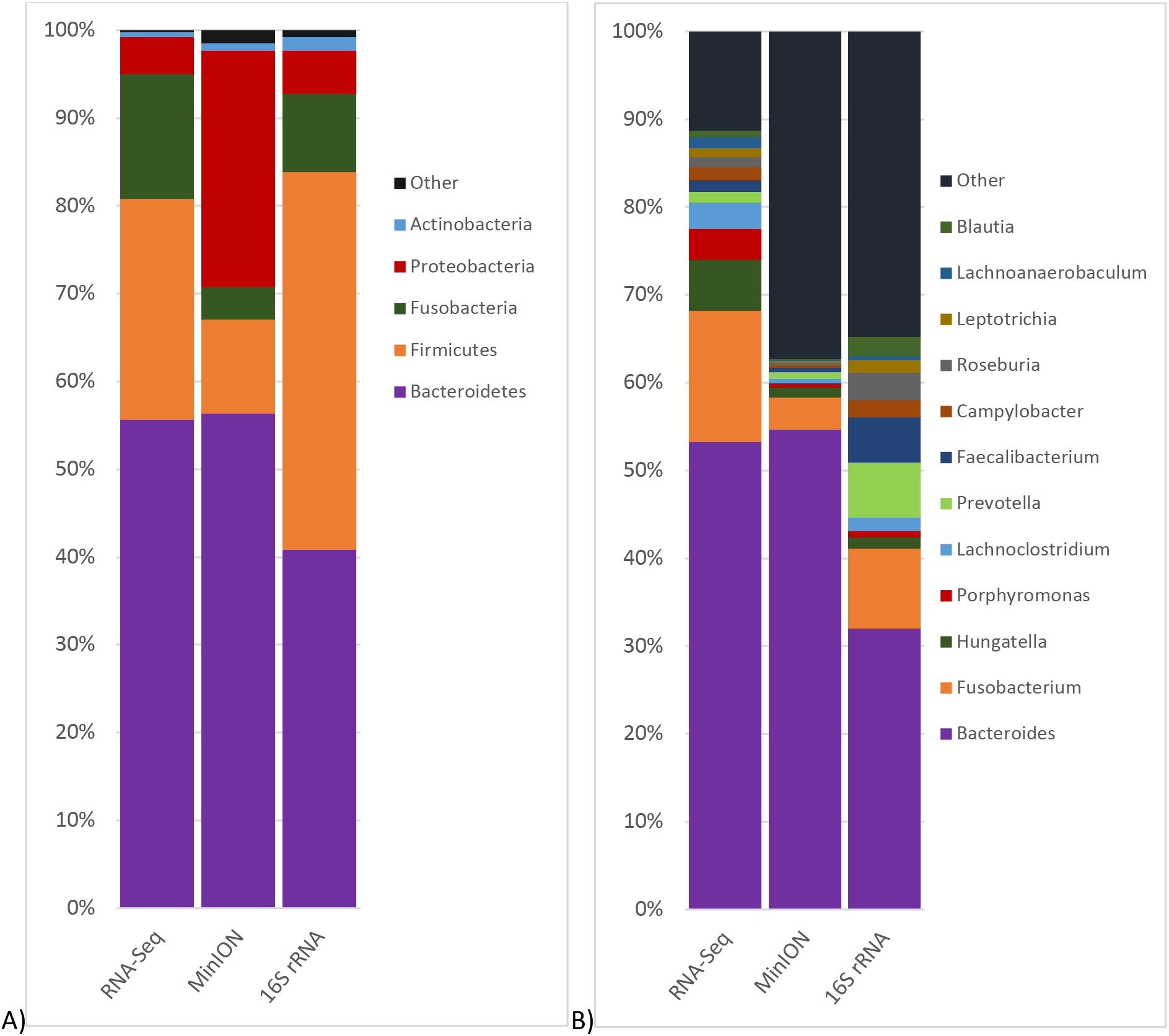
Comparison of relative abundance of between sequencing platforms at A) the phylum level and B) the genus level.

The relative abundance at the genus level also differed substantially between sequencing platforms, as shown in Figure 2B. A lower proportion of *Bacteroides* and a high proportion of *Prevotella* was detected using 16S-Seq. RNA-Seq detected a significantly larger proportion of *Fusobacterium Hungatella*, and *Porphrymonas* compared to the other platforms.

At the species level, RNA-Seq was able to detect a greater number of species than the other two platforms, while MinION-Seq could detect more than 80 species undetected by RNA-Seq or 16S sequencing. Only 689 species were detected using 16S-Seq, almost all of which were detected using at least one of the other platforms (Figure 3). The number of raw reads required to classify species is much higher for short read compared to long reads[31]. An average of 893 MinION reads was mapped to each species, while an average of 3,724 reads and 42,987 reads, mapped to a single species using 16S-Seq and RNA-Seq, respectively (Table 2).

**Table 2.** Number of different taxa detected using each sequencing platform

|  | **RNA-Seq** | **16S rRNA** | **MinION** |
| --- | --- | --- | --- |
| **Phyla Detected** | 41 | 33 | 29 |
| **Genera Detected** | 1156 | 424 | 605 |
| **Species Detected** | 3570 | 689 | 1365 |
| **Unique Phyla** | 5 | 0 | 1 |
| **Unique Genera** | 405 | 0 | 6 |
| **Unique Species** | 1944 | 9 | 81 |
| **Reads per Species** | 42,986 | 3723 | 893 |

**Figure 3.**
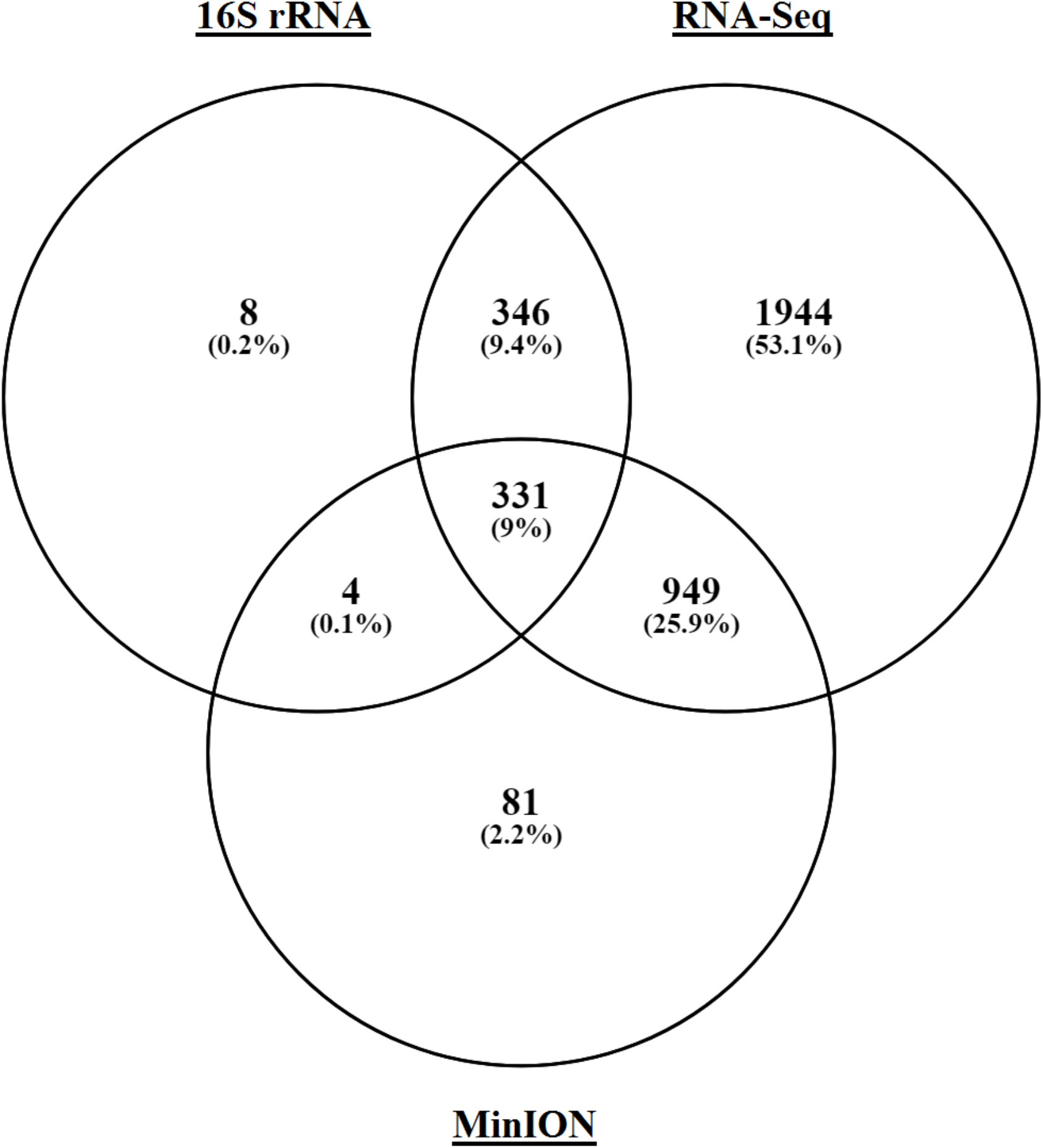
Comparison of bacterial species detection between each sequencing platform.

### Detection of colorectal cancer-associated bacterial species

Several bacterial species implicated in colorectal cancer were detected across all platforms, such as *B. fragilis* [4], *F. nucleatum* [32] and *Prevotella intermedia* [33]. Of the 81 species detected solely using MinION sequencing several were closely related *Bacillus*, *Burkholderia*, *Streptomyces* and *Psudomonas* species. Only nine species were detected using 16S-Seq that were not detected using the other two platforms (Supplementary Tables S4 and S5). RNA-Seq was able to detect 1944 species not detected by the other platforms. In addition, RNA-seq and MinION-Seq could detect several commensal species, such as *Lachnoclostridium sp* known to be involved in gut health *[34]*, that were not detected using 16S rRNA amplicon sequencing. Across all platforms, *B. fragilis* was the most abundant species, followed by *Hungatella hatheway*, *F. nucleatum*, *B. vulgatus* and *Faecalibacterium prausnitzii* (Supplementary Table S4).

Both MinION and RNA-sequencing reads were aligned to the genome of *F. nucleatum subsp. nucleatum* ATCC 25586. A single 9 kb MinION read covered a region containing four coding genes, including those coding for a putative *TetR* transcriptional regulator, involved in antibiotic resistance, and for an amino-histidine dipeptidase. Separated by four hypothetical protein coding genes, the mapped region also contained a metal-binding protein gene and manganese transport gene. RNA-Seq reads predominantly mapped to these latter two genes, with few reads mapping to the region downstream of the hypothetical protein genes (Figure 4A).

**Figure 4.**
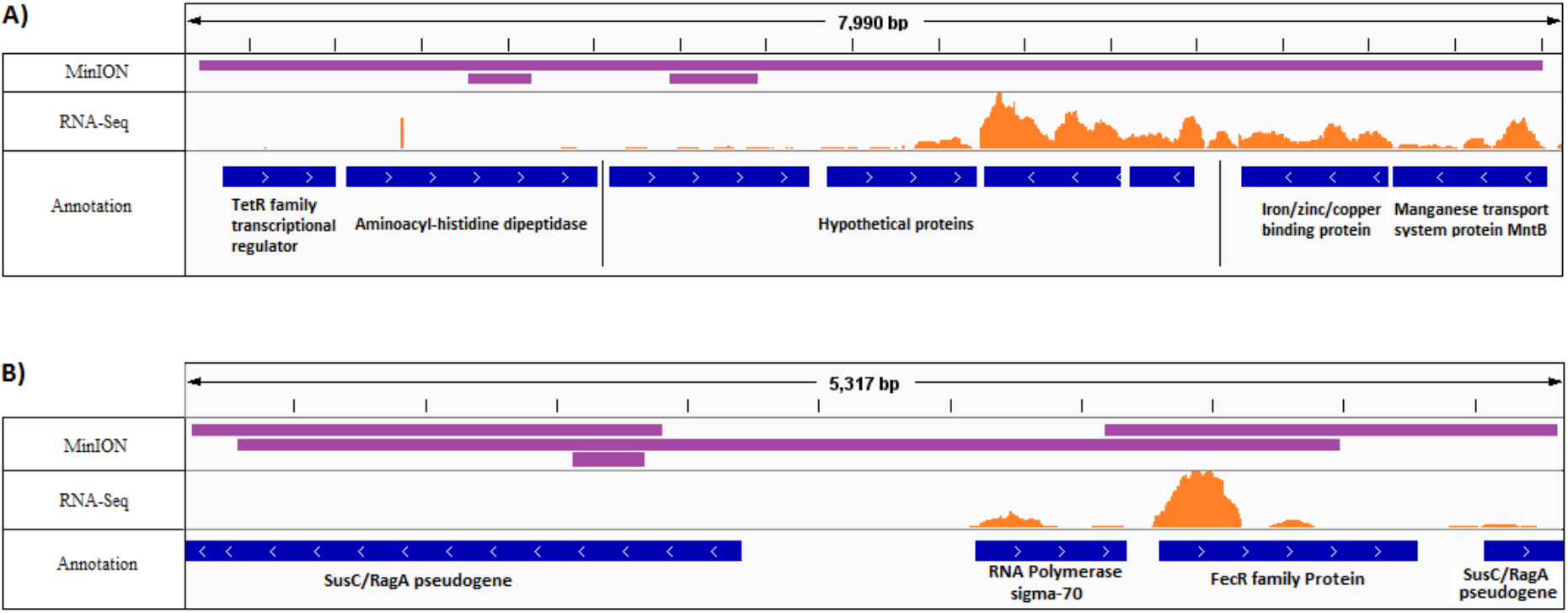
Alignment of MinION and RNA-Sequencing data to bacterial genomes of A) *F. nucleatum subsp. nucleatum* ATCC 25586 and B) *B. fragilis* Q1F2 plasmid. MinION reads are represented in purple and RNA-Sequencing reads in orange.

Additionally, sequencing reads were aligned to a conjugative plasmid of *B. fragilis* strain Q1F2 (Figure 4B). Four MinION reads, of lengths 4200, 1913, 1760, and 308bp, mapped to the plasmid to cover a 5317bp region. RNA-Seq had 75 reads align at the highest peak, aligning to a total region of 1557 bp. The majority of RNA-Seq reads mapped to the promoter region of a *FecR* iron transport gene and a corresponding sigma-70 polymerase gene.

## Discussion

Microbiome studies are increasingly being carried out on clinical samples and are expanding our knowledge of the involvement of the gut microbiome in human health and disease. Although microbiome analysis is not currently used in the clinical setting, it is likely, with the advent of microbiota-based therapeutics and prognostication, that this type of analysis will be co-opted for clinical use in the near future. The majority of published studies to date have used 16S rRNA amplicon sequencing to describe the structure of the gut microbiome. This has proven a useful tool in determining large-scale changes in the microbiome but has less utility in detecting subtle changes in microbial composition at the species level, as 16S rRNA amplicons are often too similar at the sequence level to differentiate between species [35]. Metagenomic analysis using shotgun sequencing allows for a more in-depth analysis of the microbiome to the species or even strain level, in addition to capturing mobile genetic elements and plasmids that would not be detectable using amplicon-based approaches. A recent shift towards identifying changes in microbiome function, in addition to composition, has encouraged meta-transcriptomics approaches to microbiome analysis. However, both metagenomics and meta-transcriptomics analyses generally involve short sequencing reads and necessitate high read-depth for adequate genome coverage and accurate alignment.

The Oxford Nanopore MinION is a portable sequencing device that has recently emerged as a rapid and cost-effective sequencing platform that produces long reads, and has multiple applications in clinical microbiology, such as pathogen detection [36], bacterial genome assembly [37] and plasmid and resistance gene detection [38, 39]. Specific to microbiome analysis, the utility of the MinION platform has been demonstrated in rRNA amplicon-based sequencing [40–42] and in metagenomics for outbreak analysis of bacteria [43] and viruses [44].

In this study, we compared MinION sequencing to two currently used microbiome analysis platforms, 16S rRNA amplicon sequencing and meta-transcriptomics using RNA-Sequencing. The analysis was carried out using heavily host-sequence contaminated RNA/DNA extracted from colorectal tumour tissue samples and no attempt was made to reduce the amount of host genomic material in any of the extraction methods; only 16S-Seq selected specifically for microbial DNA. For MinION-Seq, we barcoded the 11 samples and ran them as a single multiplexed analysis on a single flow-cell. As there was no amplification step involved, the numbers of reads were several orders of magnitude lower than 16S-Seq or RNA-Seq. Following a demultiplexing step, the available putatively bacterial reads per sample were as low as 7938 for some samples. One contributing factor to the low numbers of post-mapping reads was that almost half of the sequenced reads were not barcoded; which in addition to reducing the power of the analysis, may introduce a sampling bias. An enhanced library preparation protocol could increase the overall fraction of barcoded reads. Multiplexing of samples also meant that the DNA input per sample was relatively low. Sequencing a single sample or fewer multiplexed samples per flow-cell would increase the overall number of acquired reads per sample.

We used Kraken2 for rapid taxonomic identification [28] with a customised database that included many additional colorectal cancer-associated taxa for taxonomic classification of 16S-Seq, MinION-Seq and RNA-Seq data. Kraken2 uses a k-mer based algorithm that use fragmented whole genomes as the basis of taxa classification [45], without requiring large amounts of computational resources, and with increased speed compared to direct alignment of genomic sequences, such as BLAST [46].

Concordance between 16S-Seq and the other two platforms ranged from 59.5–80.5% at the phylum level but was as low as 9% at the species level. This reflects the reduced ability of 16S-Seq to differentiate between species, as low numbers of species could be identified using our sample data. Concordance between RNA-Seq and MinION-Seq was 66.7% 51.5% and 35% at the phylum, genera and species levels, respectively.

Despite the low numbers of reads acquired using MinION-Seq, more than a 1300 species could be assigned taxonomy, the majority of which were also detected in the corresponding RNA-Seq data. Specific analysis of long, high-quality MinION reads demonstrated that MinION’s longer reads can compensate for a dramatically lower number of reads, as it is possible to observe more inter-microbial genomic complexity giving a higher resolution taxonomic assignment. This was demonstrated most succinctly by Wommack et al. [31], who found that short read lengths of up to 400bp would miss 72% of the hits, that long reads of at half the read depth could detect. Comparison of the three sequencing platforms at different taxonomic levels showed that at the phylum level, all three platforms could detect a core set of common phyla, which included the most common gut-associated phyla from the Human Microbiome Project [47]. RNA-Seq was able to detect five phyla that were not detected in the MinION data, reflecting the high numbers of reads achieved with that sequencing approach, but may also be attributed to the non-synthesis-based method of sequencing MinION employs that may influence which sequences are analysed by a pore, introducing a possible selection bias. Additionally, the higher performance of RNA-Seq compared to MinION-Seq could be due to RNA-Seq transcripts being more likely to be found in databases, leading to a higher likelihood of detection and improved estimates of relative abundance [48].

Although the concordance between MinION-Seq and RNA-Seq was high at the phylum level, the overall taxonomic assignments at the species level were considerably lower, likely due to the reduced numbers of reads observed using MinION sequencing. Mapping of RNA-Seq and MinION reads to the most recent complete genome of *B. fragilis*, showed alignment to plasmids, including those that code for a putative iron transporter protein, a nutrient thought to be involved in *B. fragilis* virulence [49]. Additionally, MinION reads mapped to a region of the *F. nucleatum* genome which contains a putative antibiotic resistance gene. These results suggest that MinION-Seq can be informative for the analysis of regions of interest, such as antibiotic resistance genes and mobile elements [50]. Using a more refined protocol, it would be feasible to screen an entire plasmid using single long reads, or low read depth to detect pathogenicity and antibiotic resistance genes. The RNA-Seq platform likely did not detect the genes associated with antibiotic resistance as they were not being actively transcribed at the time. This highlights the importance of choosing the appropriate sequencing platform to suit the experimental conditions.

## Conclusions

Here, we have shown that direct microbiome sequencing of microbiomes from colorectal cancer tumour samples is feasible using the MinION platform, and gives high taxonomic concordance compared to RNA-Seq, and may be superior to 16S rRNA sequencing for taxonomic classification to the species level. We have shown that long-read sequences can compensate for low read depth for classification purposes. Our investigation has also shown the ability of MinION sequencing to discriminate between bacterial strains and detect bacterial plasmids, and shows potential as a tool for microbiome sequencing of colorectal cancers in a clinical setting.

## Acknowledgements

The authors would like to thank Helen Morrin at the Cancer Society Tissue Bank, Christchurch, and the patients involved for generously participating in this study.

## Competing interests

The authors declare that they have no competing interests

## Funding

Funding sources (Rachel Purcell): Maurice and Phyllis Paykel Trust. Gut Cancer Foundation (NZ), with support from the Hugh Green Foundation. Colorectal Surgical Society of Australia and New Zealand (CSSANZ). The funding bodies had no role in the design of the study or collection, analysis, and interpretation of data or in writing the manuscript.

## Availability of data and material

The 16S rRNA sequencing and RNA-Sequencing datasets are available in Sequence Read Archive, study SRP117763. MinION sequencing data is available from the corresponding author on reasonable request.

## Authors’ contributions

W.T. carried out MinION sequencing, bioinformatics analysis and preparation of figures and contributed to manuscript writing. J.P was involved in study design, bioinformatics and data analysis, and manuscript preparation. A.M was involved in MinION sample preparation and sequencing. S.S. was involved in study design and bioinformatics analysis. F.F. was involved in study design and clinical aspects of the study. R.P. carried out sequencing preparation of tumour samples, MinION sequencing and was a major contributor to study design and manuscript writing. All authors read and approved the final manuscript.

## References

1. Siegel, R.L., K.D. Miller, and A. Jemal, Cancer statistics, 2018. CA Cancer J Clin, 2018. 68(1): p. 7–30.

2. Ferlay, J., et al., Cancer incidence and mortality worldwide: Sources, methods and major patterns in GLOBOCAN 2012. International Journal of Cancer, 2014. 136(5): p. E359–E386.

3. Navarro, M., et al., Colorectal cancer population screening programs worldwide in 2016: An update. World Journal of Gastroenterology, 2017. 23(20): p. 3632–3642.

4. Purcell, R.V., et al., Colonization with enterotoxigenic Bacteroides fragilis is associated with early-stage colorectal neoplasia. PLoS One, 2017. 12(2): p. e0171602.

5. Schulz, T.R., et al., Using Stool Antigen to Screen for Helicobacter pylori in Immigrants and Refugees from High Prevalence Countries Is Relatively Cost Effective in Reducing the Burden of Gastric Cancer and Peptic Ulceration. PLoS ONE, 2014. 9(9): p. e108610.

6. de Lecea, M.G.M. and M. Rossbach, Translational genomics in personalized medicine – scientific challenges en route to clinical practice. The HUGO Journal, 2012. 6(1): p. 2.

7. Quail, M.A., et al., A tale of three next generation sequencing platforms: comparison of Ion Torrent, Pacific Biosciences and Illumina MiSeq sequencers. BMC Genomics, 2012. 13(1): p. 341.

8. Scholz, M.B., C.-C. Lo, and P.S.G. Chain, Next generation sequencing and bioinformatic bottlenecks: the current state of metagenomic data analysis. Current Opinion in Biotechnology, 2012. 23(1): p. 9–15.

9. Větrovský, T. and P. Baldrian, The Variability of the 16S rRNA Gene in Bacterial Genomes and Its Consequences for Bacterial Community Analyses. PLOS ONE, 2013. 8(2): p. e57923.

10. Osman, M.A., et al., 16S rRNA gene sequencing for deciphering the colorectal cancer gut microbiome: Current protocols and workflows. Frontiers in Microbiology, 2018. 9(APR).

11. Poretsky, R., et al., Strengths and Limitations of 16S rRNA Gene Amplicon Sequencing in Revealing Temporal Microbial Community Dynamics. PLOS ONE, 2014. 9(4): p. e93827.

12. Ravi, A., et al., Transmission and persistence of IncF conjugative plasmids in the gut microbiota of full-term infants. FEMS Microbiol Ecol, 2018. 94(1).

13. Szabo, M., et al., Characterization of Two Multidrug-Resistant IncA/C Plasmids from the 1960s by Using the MinION Sequencer Device. Antimicrob Agents Chemother, 2016. 60(11): p. 6780–6786.

14. Giannoukos, G., et al., Efficient and robust RNA-seq process for cultured bacteria and complex community transcriptomes. Genome Biol, 2012. 13(3): p. R23.

15. Yan, Y.-W., et al., Modified RNA-seq method for microbial community and diversity analysis using rRNA in different types of environmental samples. PLOS ONE, 2017. 12(10): p. e0186161.

16. Krizanovic, K., et al., Evaluation of tools for long read RNA-seq splice-aware alignment. Bioinformatics, 2018. 34(5): p. 748–754.

17. Jain, M., et al., The Oxford Nanopore MinION: delivery of nanopore sequencing to the genomics community. Genome Biology, 2016. 17: p. 239.

18. Purcell, R.V., et al., Distinct gut microbiome patterns associate with consensus molecular subtypes of colorectal cancer. Sci Rep, 2017. 7(1): p. 11590.

19. Purcell, R.V.V., M.; Biggs, P. J.; Schmeier, S.; Frizelle, F. A., NCBI Sequence Read Archive SRP117763. 2017.

20. Magoc, T. and S.L. Salzberg, FLASH: fast length adjustment of short reads to improve genome assemblies. Bioinformatics, 2011. 27(21): p. 2957–63.

21. Cox, M.P., D.A. Peterson, and P.J. Biggs, SolexaQA: At-a-glance quality assessment of Illumina second-generation sequencing data. BMC Bioinformatics, 2010. 11: p. 485.

22. Caporaso, J.G., et al., QIIME allows analysis of high-throughput community sequencing data. Nat Methods, 2010. 7(5): p. 335–6.

23. Dobin, A., et al., STAR: ultrafast universal RNA-seq aligner. Bioinformatics, 2013. 29(1): p. 15–21.

24. Li, H., et al., The Sequence Alignment/Map format and SAMtools. Bioinformatics, 2009. 25(16): p. 2078–9.

25. Quinlan, A.R. and I.M. Hall, BEDTools: a flexible suite of utilities for comparing genomic features. Bioinformatics, 2010. 26(6): p. 841–842.

26. De Coster, W., et al., NanoPack: visualizing and processing long-read sequencing data. Bioinformatics, 2018. 34(15): p. 2666–2669.

27. Li, H., Minimap2: pairwise alignment for nucleotide sequences. Bioinformatics, 2018: p. bty191–bty191.

28. Wood, D.E. and S.L. Salzberg, Kraken: ultrafast metagenomic sequence classification using exact alignments. Genome Biol, 2014. 15(3): p. R46.

29. Breitwieser, F.P. and S.L. Salzberg, Pavian: Interactive analysis of metagenomics data for microbiomics and pathogen identification. bioRxiv, 2016: p. 084715.

30. Robinson, J.T., et al., Integrative Genomics Viewer. Nature biotechnology, 2011. 29(1): p. 24–26.

31. Wommack, K.E., J. Bhavsar, and J. Ravel, Metagenomics: Read Length Matters. Applied and Environmental Microbiology, 2008. 74(5): p. 1453–1463.

32. Castellarin, M., et al., Fusobacterium nucleatum infection is prevalent in human colorectal carcinoma. Genome Research, 2012. 22(2): p. 299–306.

33. Flynn, K.J., N.T. Baxter, and P.D. Schloss, Metabolic and Community Synergy of Oral Bacteria in Colorectal Cancer. mSphere, 2016. 1(3).

34. Biddle, A., et al., Untangling the Genetic Basis of Fibrolytic Specialization by Lachnospiraceae and Ruminococcaceae in Diverse Gut Communities. Diversity, 2013. 5(3).

35. Stary, L., et al., Are we any closer to screening for colorectal cancer using microbial markers?A critical review. Biomed Pap Med Fac Univ Palacky Olomouc Czech Repub, 2017. 161(4): p. 333–338.

36. Gong, L., et al., Culture-independent analysis of liver abscess using nanopore sequencing. PLoS One, 2018. 13(1): p. e0190853.

37. Wick, R.R., et al., Completing bacterial genome assemblies with multiplex MinION sequencing. Microb Genom, 2017. 3(10): p. e000132.

38. Lemon, J.K., et al., Rapid Nanopore Sequencing of Plasmids and Resistance Gene Detection in Clinical Isolates. J Clin Microbiol, 2017. 55(12): p. 3530–3543.

39. van der Helm, E., et al., Rapid resistome mapping using nanopore sequencing. Nucleic Acids Res, 2017. 45(8): p. e61.

40. Kerkhof, L.J., et al., Profiling bacterial communities by MinION sequencing of ribosomal operons. Microbiome, 2017. 5(1): p. 116.

41. Mitsuhashi, S., et al., A portable system for rapid bacterial composition analysis using a nanopore-based sequencer and laptop computer. Sci Rep, 2017. 7(1): p. 5657.

42. Benitez-Paez, A. and Y. Sanz, Multi-locus and long amplicon sequencing approach to study microbial diversity at species level using the MinION portable nanopore sequencer. Gigascience, 2017. 6(7): p. 1–12.

43. Walter, M.C., et al., MinION as part of a biomedical rapidly deployable laboratory. J Biotechnol, 2017. 250: p. 16–22.

44. Batovska, J., et al., Metagenomic arbovirus detection using MinION nanopore sequencing. J Virol Methods, 2017. 249: p. 79–84.

45. Carvalho, A.B., E.G. Dupim, and G. Goldstein, Improved assembly of noisy long reads by k-mer validation. Genome Res, 2016. 26(12): p. 1710–1720.

46. Camacho, C., et al., BLAST+: architecture and applications. BMC Bioinformatics, 2009. 10: p. 421.

47. Group, N.H.W., et al., The NIH Human Microbiome Project. Genome Res, 2009. 19(12): p. 2317–23.

48. Cottier, F., et al., Advantages of meta-total RNA sequencing (MeTRS) over shotgun metagenomics and amplicon-based sequencing in the profiling of complex microbial communities. npj Biofilms and Microbiomes, 2018. 4(1): p. 2.

49. Robertson, K.P., et al., Characterization of Bacteroides fragilis hemolysins and regulation and synergistic interactions of HlyA and HlyB. Infect Immun, 2006. 74(4): p. 2304–16.

50. Garalde, D.R., et al., Highly parallel direct RNA sequencing on an array of nanopores. Nature Methods, 2018. 15: p. 201.

